# Genomic and Transcriptomic Profiling of Phoenix Colonies

**DOI:** 10.1101/2022.03.05.483082

**Authors:** Devin Sindeldecker, Matthew Dunn, Aubree Zimmer, Matthew Anderson, Juan Alfonzo, Paul Stoodley

## Abstract

*Pseudomonas aeruginosa* is a Gram-negative bacterium responsible for numerous human infections. Previously, novel antibiotic tolerant variants known as phoenix colonies as well as variants similar to viable but non-culturable (VBNC) colonies were identified in response to high concentrations of aminoglycosides. In this study, the mechanisms behind phoenix colony and VBNC-like colony emergence were further explored using both whole genome sequencing and RNA sequencing. Phoenix colonies were found to have a single nucleotide polymorphism (SNP) in the PA4673 gene, which is predicted to encode a GTP-binding protein. No SNPs were identified within VBNC-like colonies compared to the founder population. RNA sequencing did not detect change in expression of PA4673 but revealed multiple differentially expressed genes that may play a role in phoenix colony emergence. One of these differentially expressed genes, PA3626, encodes for a tRNA pseudouridine synthase which when knocked out led to a complete lack of phoenix colonies. Although not immediately clear whether the identified genes in this study may have interactions which have not yet been recognized, they may contribute to the understanding of how phoenix colonies are able to emerge and survive in the presence of antibiotic exposure.

## Introduction

*Pseudomonas aeruginosa* is a Gram-negative bacterium found throughout the natural environment. As an opportunistic pathogen, it is most commonly associated with cystic fibrosis (CF) infections but can also reside in chronic wounds and post-surgical site infections (1–3). *P. aeruginosa* utilizes the formation of biofilms, persister cells, and development of multidrug resistance mechanisms to evade killing by antimicrobial agents (4–7). In addition to these antimicrobial tolerance and resistance mechanisms, *P. aeruginosa* has been found to rapidly adapt to its environment within the context of infection, leading to concerns of reduced effectiveness of antimicrobial agents in eradicating *P. aeruginosa* (8).

In a previous study, our lab identified a novel, aminoglycoside tolerant phenotype of *P. aeruginosa*, which we have termed phoenix colonies. Phoenix colonies are able to survive and thrive in an antibiotic laden environment. However, once removed from the initial antibiotic environment, phoenix colonies revert to a wild-type level of antibiotic susceptibility (9). Phoenix colonies differ from traditional tolerance in that they are able to maintain high levels of metabolic activity throughout antibiotic exposure, whereas traditional tolerance is typically the result of slow growth or a decrease in metabolism (10). In the same study, we identified an additional phenotype which was similar to the viable but non-culturable colonies (VBNCs, (11)) phenotype, however, these “VBNC-like” colonies were able to grow in the initial antibiotic environment but were unable to be cultured otherwise, including cultures containing the same antibiotic (9). While the significance of these phenotypes in a clinical setting is currently unknown, it is possible that phoenix colonies or VBNC-like colonies could lead to chronic or recurrent infections, similar to persister cells (7). Furthermore, the molecular and genetic mechanisms leading to phoenix colony and VBNC-like emergence are unknown. Understanding the genetic alterations in these cells or alterations to gene expression may allow for better preventative or treatment options in the management of chronic or recurrent infections.

In this study, we used whole genome sequencing (WGS) of phoenix colonies and VBNC-like colonies to identify a single nucleotide polymorphism (SNP) which may be associated with the emergence of phoenix colonies in the presence of high concentrations of aminoglycosides. Additionally, RNA sequencing was performed on phoenix colonies as well as VBNC-like colonies and identified differentially expressed genes (DEGs) which could account for the antibiotic adaptation phenotypes portrayed by both phoenix colonies and VBNC-like colonies.

## Results

### Phoenix Colonies Contain a SNP in PA4673

As shown previously (9), antibiotic resistant and tolerant variant colonies were cultured by exposing a *P. aeruginosa* PAO1 biofilm lawn to high concentrations of tobramycin (Figure 1). Genomic DNA was isolated from phoenix colonies, VBNC-like colonies, and the founder population of *P. aeruginosa* PAO1. Whole genome sequencing (WGS) was performed using a Nanopore MinION Sequencer, and the reads were mapped to the PAO1 reference genome. Mapped reads were analyzed for the presence of SNPs as well as insertions or deletions (indels). Only the phoenix colony sample encoded any SNPs compared to the PAO1 reference genome and no samples (phoenix colony, VBNC-like colony, or control colony) contained indels. The single guanine to thymine (GCF_000006765.1:c.353G>T) variant occurred within the PA4673 gene, a hypothetical cytosolic protein, to produce a serine to isoleucine (S118I) amino acid change.

**Figure 1.**
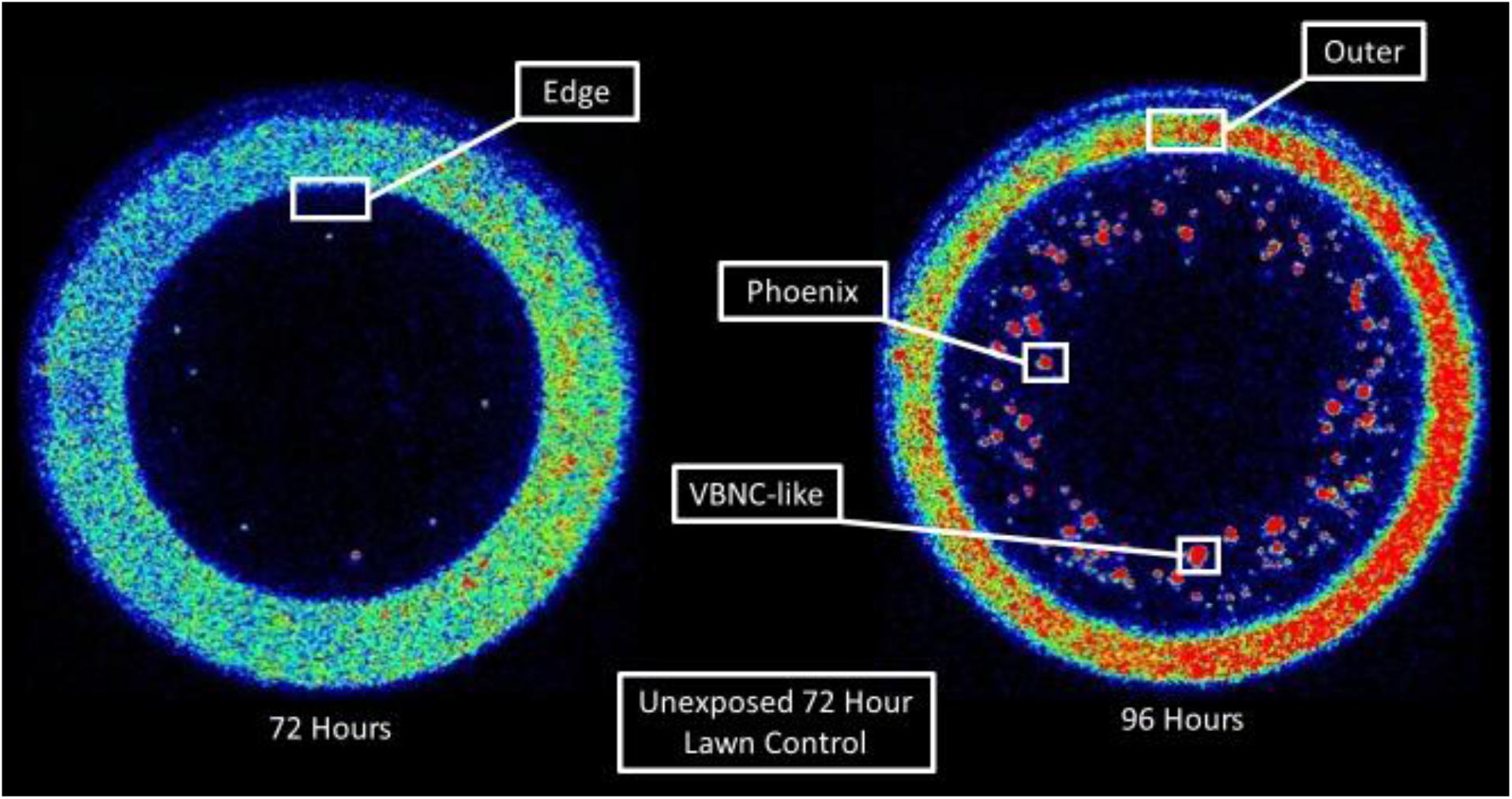
Isolation of *P. aeruginosa* colony variants for assaying the genome and gene expression. *In vitro* imaging system (IVIS) images of representative phoenix colony plates show areas of bacterial sampling. Seventy-two hours post tobramycin bead placement, samples were taken from the edge of the zone of clearance (ZOC) where phoenix colonies would be expected to arise the following day after continued incubation. Ninety-six hours post tobramycin bead placement, samples were taken from phoenix colonies, VBNC-like colonies, and the outer background lawn. Additionally, samples were taken from bacterial lawns grown for 72 hours that had not been exposed to antibiotics to be used as comparative controls. Red indicates high levels of metabolic activity, blue indicates low levels of metabolic activity, black indicates no metabolic activity.

### PA4673 is Predicted to Encode a GTP-binding Protein

To determine the impact of the S118I polymorphism in PA4673, *de novo* protein modeling was performed using Phyre2 (12). The structural models of both the wild-type and mutant proteins show a change in the secondary structure of the mutant protein, specifically, the formation of a predicted β sheet at the mutation site, which is located in an exposed face of the protein and is also predicted to have an altered hydrophobicity in the mutated region (Figure 2, Figure 3). This modification may lead to alteration of a binding site or protein function. Identification of functional domains in both PA4673 isoforms predicted domains between AA001 and AA363, indicating they likely function as GTP-binding proteins as they align with 100% confidence to the YchF GTP-binding protein of *Schizosaccharomyces pombe* (13, 14).

**Figure 2.**
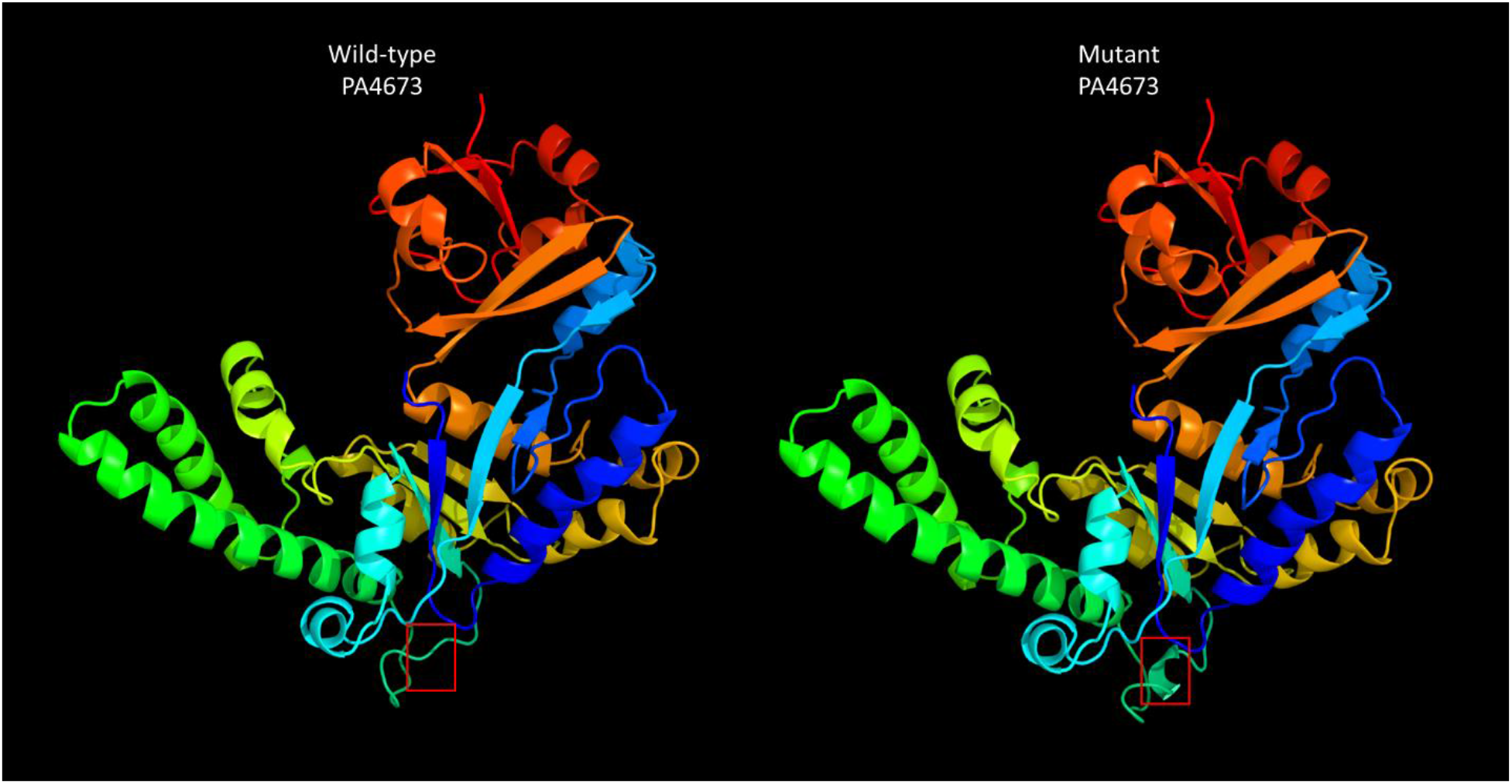
Protein structure of wild-type and mutant PA4673. Protein structures were derived from the amino acid sequences of wild-type and mutant PA4673 using Phyre2. The red box indicates the structural change caused by the SNP. Functional domain identification in both PA4673 isoforms indicates that they likely function as GTP-binding proteins. As the SNP causes an amino acid change in the exterior portion of the predicted protein structure, it may lead to alteration of a binding site or protein function.

**Figure 3.**
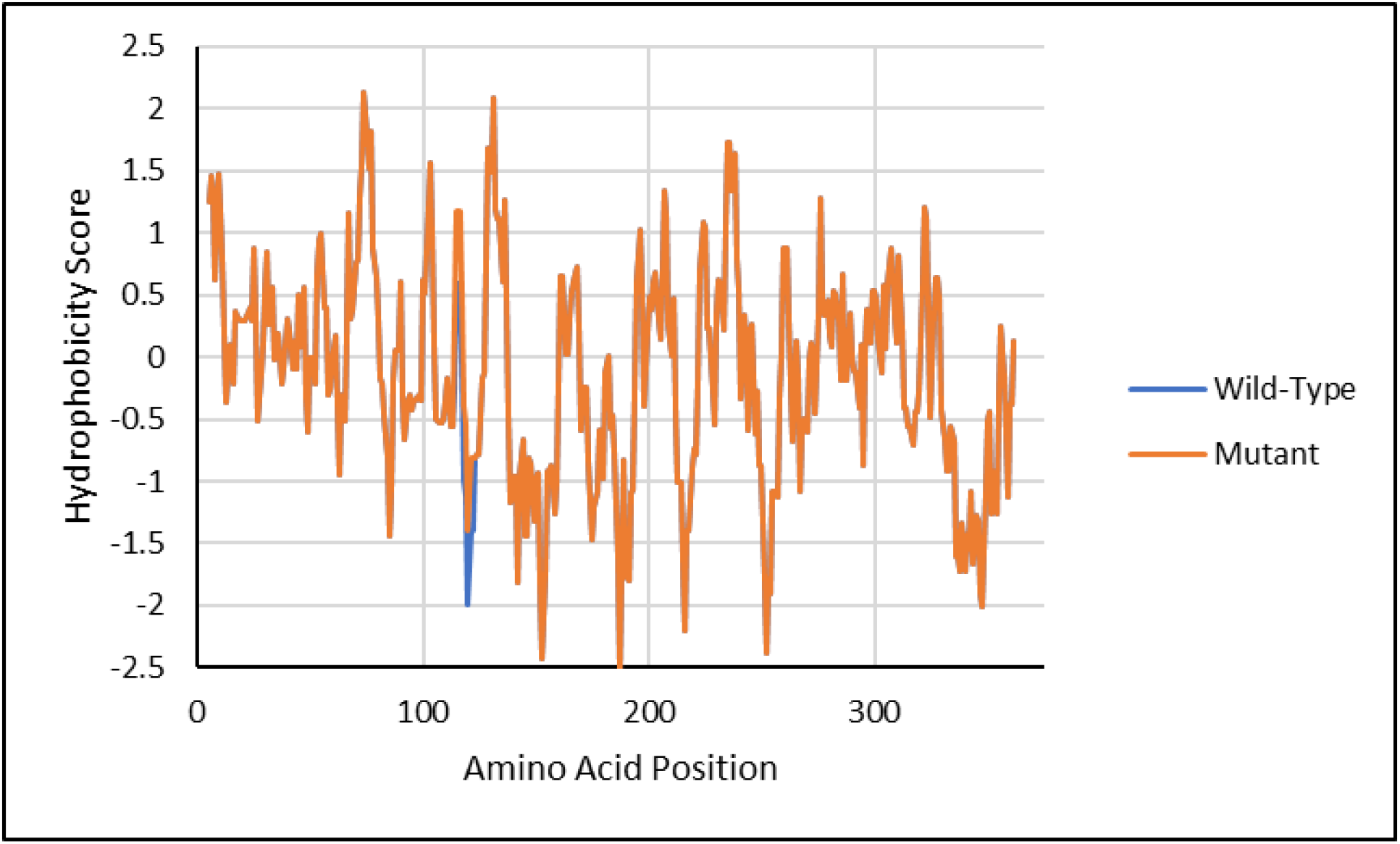
Hydrophobicity of wild-type and mutant PA4673 proteins. Amino acid sequences for wild-type and the SNP containing mutant PA4673 proteins were submitted to the ExPASy server for amino acid hydrophobicity score prediction. A change in hydrophobicity is noted in the region of the mutant amino acid which may alter binding of the protein to substrates.

### DEGs were Identified for Phoenix Colonies and VBNC-like Colonies

Due to the lack of clear genetic candidates for promoting alternative colony states, a transcriptomic approach was utilized to define gene regulation changes that are associated with each alternative colony type. RNA was isolated from samples at the edge of the ZOC at 72 hours, from multiple colony populations at 96 hours (phoenix colonies, VBNC-like colonies, and the outer background lawn), and from a 72 hour unexposed lawn as a control, and prepared for Illumina-based sequencing (Figure 1).

Differential gene expression of either phoenix colonies or VBNC-like colonies was performed to define transcriptional profiles that distinguish each phenotype. First, a multi-dimensional scaling (MDS) plot was generated using expression of all genes to examine the relationships between all samples (Figure 4). This plot shows compact clustering among biological replicates of each colony phenotype with the exception of 96-hour outer-edge populations. While the edge of ZOC group, control, and outer background lawn samples clustered on their own. The phoenix colonies and VBNC-like colonies overlap heavily, indicating highly similar transcriptomic profiles. Differential gene expression between the control lawn and phoenix colonies, as well as the control lawn and VBNC-like colonies, identified 63 and 90 genes with significantly different transcript abundance, respectively (>2x fold change, corrected p value<0.05, Figure 5). These genes were split with 40 down-regulated and 23 up-regulated genes when comparing phoenix colonies to the control lawn, and 56 down-regulated and 34 up-regulated genes for VBNC-like colonies compared to the control (Table 1, Table 2). The DEGs were submitted for DAVID, KEGG, and Gene Ontology analysis, but no significant correlation was identified.

**Table 1.** Statistically significant DEGs of phoenix colonies. Phoenix colony transcripts were compared to the control lawn to identify genes which were significantly up or down regulated. Red indicates genes which were significantly upregulated.

| Gene | Fold Change | q-value | Protein Product Annotation |
| --- | --- | --- | --- |
| PA4746 | 2.63E+01 | 4.79E-02 | hypothetical protein PA4746 |
| PA3476 | 1.85E+01 | 4.62E-02 | acyl-homoserine-lactone synthase |
| PA3806 | 3.14E+01 | 4.50E-02 | dual-specificity RNA methyltransferase RlmN |
| PA2561 | 4.15E-02 | 4.43E-02 | methyl-accepting chemotaxis protein CtpH |
| PA3699 | 4.27E+01 | 4.40E-02 | transcriptional regulator |
| PA2213 | 3.28E+01 | 4.31E-02 | porin |
| PA3626 | 3.78E-02 | 4.23E-02 | tRNA pseudouridine synthase D |
| PA3385 | 2.49E+01 | 4.23E-02 | alginate and motility regulator Z |
| PA1825 | 2.94E-02 | 4.17E-02 | hypothetical protein PA1825 |
| PA2751 | 7.81E-02 | 4.14E-02 | hypothetical protein PA2751 |
| PA3080 | 3.77E-02 | 4.00E-02 | hypothetical protein PA3080 |
| PA3719 | 2.56E+01 | 3.91E-02 | MexR antirepressor ArmR |
| PA1771 | 6.45E-02 | 3.80E-02 | hydrolase |
| PA2819.3 | 4.88E+01 | 3.79E-02 | tRNA-Glu |
| PA1673 | 4.22E+01 | 3.63E-02 | bacteriohemerythrin |
| PA4529 | 5.78E+01 | 3.63E-02 | dephospho-CoA kinase |
| PA4746.2 | 1.42E+02 | 3.63E-02 | tRNA-Leu |
| PA1037 | 2.89E-02 | 3.38E-02 | hypothetical protein PA1037 |
| PA2843 | 1.79E-02 | 2.58E-02 | aldolase |
| PA2738 | 2.66E-02 | 2.21E-02 | integration host factor subunit alpha |
| PA0107 | 3.66E-02 | 2.21E-02 | cytochrome C oxidase assembly protein |
| PA1278 | 1.68E+01 | 2.21E-02 | adenosylcobinamide kinase/adenosylcobinamide-phosphate guanylyltransferase |
| PA5508 | 3.79E+01 | 2.21E-02 | glutamine synthetase |
| PA4746.1 | 8.10E+01 | 2.21E-02 | tRNA-Met |
| PA2948 | 1.82E-02 | 2.08E-02 | precorrin-4 C(11)-methyltransferase |
| PA4511 | 4.68E-03 | 1.92E-02 | hypothetical protein PA4511 |
| PA2010 | 3.66E+01 | 1.92E-02 | transcriptional regulator |
| PA3973 | 3.87E-02 | 1.56E-02 | transcriptional regulator |
| PA1425 | 1.17E-02 | 9.79E-03 | ABC transporter ATP-binding protein |
| PA0846 | 2.50E-02 | 9.79E-03 | sulfate transporter CysZ |
| PA3399 | 1.96E+02 | 7.55E-03 | hypothetical protein PA3399 |
| PA2573 | 6.43E-03 | 7.23E-03 | chemotaxis transducer |
| PA4591 | 1.78E-02 | 6.24E-03 | hypothetical protein PA4591 |
| PA1833 | 1.98E-02 | 6.24E-03 | oxidoreductase |
| PA0298 | 4.59E-02 | 5.93E-03 | glutamine synthetase |
| PA0760 | 8.50E+01 | 5.93E-03 | hypothetical protein PA0760 |
| PA1409 | 5.86E-03 | 5.91E-03 | acetylpolyamine aminohydrolase |
| PA3611 | 1.84E-02 | 5.91E-03 | hypothetical protein PA3611 |
| PA5210 | 1.29E-02 | 5.90E-03 | secretion pathway ATPase |
| PA3464 | 2.87E-02 | 5.55E-03 | hypothetical protein PA3464 |
| PA3115 | 5.27E+02 | 5.55E-03 | motility protein FimV |
| PA2482 | 6.50E-03 | 4.87E-03 | cytochrome C |
| PA3708 | 7.31E+01 | 4.87E-03 | chemotaxis transducer |
| PA4362 | 1.31E-02 | 4.15E-03 | hypothetical protein PA4362 |
| PA1146 | 9.04E-03 | 3.31E-03 | iron-containing alcohol dehydrogenase |
| PA5330 | 1.24E-02 | 3.31E-03 | hypothetical protein PA5330 |
| PA0103a | 1.66E-02 | 3.31E-03 | hypothetical protein PA0103a |
| PA0727 | 7.69E+01 | 3.31E-03 | hypothetical protein PA0727 |
| PA2139 | 1.67E-02 | 2.64E-03 | pseudogene |
| PA3405 | 1.50E-02 | 1.94E-03 | metalloprotease secretion protein |
| PA3718 | 9.32E+01 | 1.94E-03 | major facilitator superfamily transporter |
| PA4745 | 2.43E+02 | 1.30E-03 | transcription elongation factor NusA |
| PA3610 | 1.39E-02 | 4.19E-04 | polyamine ABC transporter substrate-binding protein<br>PotD |
| PA0608 | 2.43E+02 | 4.19E-04 | phosphoglycolate phosphatase |
| PA2503 | 1.22E-02 | 1.93E-04 | hypothetical protein PA2503 |
| PA4492 | 7.26E-03 | 1.02E-04 | hypothetical protein PA4492 |
| PA2926 | 2.67E-03 | 9.37E-05 | histidine ABC transporter ATP-binding protein |
| PA4713 | 5.33E-04 | 5.06E-05 | hypothetical protein PA4713 |
| PA3355 | 1.14E-03 | 2.93E-05 | hypothetical protein PA3355 |
| PA4712 | 6.37E-04 | 3.33E-06 | hypothetical protein PA4712 |
| PA3038 | 3.93E-04 | 1.89E-08 | porin |
| PA3609 | 3.59E-05 | 1.10E-09 | polyamine ABC transporter permease PotC |
| PA0526 | 1.68E-06 | 6.45E-20 | hypothetical protein PA0526 |

**Table 2.** Statistically significant DEGs of VBNC-like colonies. VBNC-like colony transcripts were compared to the control lawn to identify genes which were significantly up or down regulated. Red indicates genes which were significantly upregulated and blue indicates genes which were significantly downregulated.

| Gene | Fold Change | q-value | Protein Product Annotation |
| --- | --- | --- | --- |
| PA0526 | 9.46E-07 | 2.80E-21 | hypothetical protein PA0526 |
| PA3609 | 1.74E-05 | 6.22E-11 | polyamine ABC transporter permease PotC |
| PA3038 | 2.10E-04 | 8.83E-10 | porin |
| PA3355 | 1.32E-04 | 4.61E-09 | hypothetical protein PA3355 |
| PA4712 | 8.10E-04 | 6.81E-06 | hypothetical protein PA4712 |
| PA2926 | 2.12E-03 | 4.29E-05 | histidine ABC transporter ATP-binding protein HisP |
| PA2503 | 9.06E-03 | 5.02E-05 | hypothetical protein PA2503 |
| PA3464 | 1.03E-02 | 1.24E-04 | hypothetical protein PA3464 |
| PA4591 | 6.02E-03 | 2.75E-04 | hypothetical protein PA4591 |
| PA4713 | 1.16E-03 | 3.30E-04 | hypothetical protein PA4713 |
| PA1146 | 5.03E-03 | 5.38E-04 | iron-containing alcohol dehydrogenase |
| PA3610 | 1.56E-02 | 6.08E-04 | polyamine ABC transporter substrate-binding protein<br>PotD |
| PA1833 | 8.90E-03 | 6.08E-04 | oxidoreductase |
| PA4724.1 | 6.57E-03 | 6.27E-04 | hypothetical protein PA4724.1 |
| PA4492 | 1.21E-02 | 6.32E-04 | hypothetical protein PA4492 |
| PA5210 | 6.14E-03 | 6.32E-04 | secretion pathway ATPase |
| PA2561 | 9.94E-03 | 7.12E-04 | methyl-accepting chemotaxis protein CtpH |
| PA1083 | 9.13E+01 | 7.33E-04 | flagellar basal body L-ring protein |
| PA3080 | 1.06E-02 | 8.81E-04 | hypothetical protein PA3080 |
| PA2573 | 2.71E-03 | 8.81E-04 | chemotaxis transducer |
| PA2843 | 5.00E-03 | 9.36E-04 | aldolase |
| PA2482 | 3.89E-03 | 9.36E-04 | cytochrome C |
| PA3405 | 1.54E-02 | 1.25E-03 | metalloprotease secretion protein |
| PA0103a | 1.50E-02 | 1.57E-03 | hypothetical protein PA0103a |
| PA1425 | 5.96E-03 | 1.57E-03 | ABC transporter ATP-binding protein |
| PA2738 | 1.16E-02 | 1.91E-03 | integration host factor subunit alpha |
| PA3611 | 1.38E-02 | 1.97E-03 | hypothetical protein PA3611 |
| PA5330 | 1.20E-02 | 2.03E-03 | hypothetical protein PA5330 |
| PA0727 | 7.05E+01 | 3.01E-03 | hypothetical protein PA0727 |
| PA0939 | 3.48E+01 | 3.14E-03 | hypothetical protein PA0939 |
| PA1409 | 5.63E-03 | 3.86E-03 | acetylpolyamine aminohydrolase |
| PA2948 | 1.01E-02 | 4.04E-03 | precorrin-4 C(11)-methyltransferase |
| PA2565 | 2.96E-02 | 4.16E-03 | hypothetical protein PA2565 |
| PA1825 | 1.25E-02 | 4.26E-03 | hypothetical protein PA1825 |
| PA0846 | 2.21E-02 | 5.67E-03 | sulfate transporter CysZ |
| PA4362 | 1.62E-02 | 5.67E-03 | hypothetical protein PA4362 |
| PA3718 | 5.46E+01 | 5.70E-03 | major facilitator superfamily transporter |
| PA1673 | 9.02E+01 | 6.92E-03 | bacteriohemerythrin |
| PA2167 | 2.77E+01 | 1.19E-02 | hypothetical protein PA2167 |
| PA0107 | 3.18E-02 | 1.23E-02 | cytochrome C oxidase assembly protein |
| PA3406 | 2.88E-02 | 1.23E-02 | transporter HasD |
| PA3626 | 2.39E-02 | 1.23E-02 | tRNA pseudouridine synthase D |
| PA3973 | 3.86E-02 | 1.31E-02 | transcriptional regulator |
| PA2613 | 3.23E-02 | 1.31E-02 | recombination factor protein RarA |
| PA1418 | 1.65E+01 | 1.47E-02 | sodium:solute symport protein |
| PA2139 | 3.23E-02 | 1.47E-02 | pseudogene |
| PA2819.3 | 7.20E+01 | 1.48E-02 | tRNA-Glu |
| PA2942 | 5.81E+01 | 1.85E-02 | magnesium chelatase |
| PA1590 | 2.94E+01 | 1.85E-02 | branched-chain amino acid transporter |
| PA1037 | 2.44E-02 | 1.85E-02 | hypothetical protein PA1037 |
| PA0039 | 1.94E-02 | 1.94E-02 | hypothetical protein PA0039 |
| PA4510 | 4.48E-02 | 2.00E-02 | hypothetical protein PA4510 |
| PA0938 | 1.85E+01 | 2.07E-02 | hypothetical protein PA0938 |
| PA0263.1 | 1.83E+01 | 2.07E-02 | tRNA-Arg |
| PA4127 | 2.06E-02 | 2.09E-02 | 2-oxo-hepta-3-ene-1,7-dioic acid hydratase |
| PA4370 | 6.07E+01 | 2.16E-02 | insulin-cleaving metalloproteinase outer membrane protein |
| PA4746 | 3.54E+01 | 2.16E-02 | hypothetical protein PA4746 |
| PA5353 | 7.62E-03 | 2.40E-02 | glycolate oxidase iron-sulfur subunit |
| PA3115 | 1.67E+02 | 2.49E-02 | motility protein FimV |
| PA3246 | 8.96E+01 | 2.49E-02 | pseudouridine synthase |
| PA4746.1 | 6.77E+01 | 2.49E-02 | tRNA-Met |
| PA0884 | 1.78E+01 | 2.49E-02 | C4-dicarboxylate-binding protein |
| PA0650 | 7.72E-02 | 2.49E-02 | anthranilate phosphoribosyltransferase |
| PA3719 | 2.70E+01 | 2.53E-02 | MexR antirepressor ArmR |
| PA3608 | 2.04E+01 | 2.53E-02 | polyamine ABC transporter permease PotB |
| PA0298 | 7.48E-02 | 2.53E-02 | glutamine synthetase |
| PA3711 | 5.45E-02 | 2.53E-02 | transcriptional regulator |
| PA4529 | 5.39E+01 | 2.89E-02 | dephospho-CoA kinase |
| PA3636 | 7.44E-02 | 2.89E-02 | 2-dehydro-3-deoxyphosphooctonate aldolase |
| PA1154 | 3.41E-02 | 2.89E-02 | hypothetical protein PA1154 |
| PA5506 | 2.40E-02 | 2.89E-02 | hypothetical protein PA5506 |
| PA3385 | 2.69E+01 | 2.92E-02 | alginate and motility regulator Z |
| PA5367 | 6.56E-02 | 2.92E-02 | phosphate ABC transporter permease |
| PA4652 | 3.76E-02 | 2.92E-02 | hypothetical protein PA4652 |
| PA4746.2 | 1.16E+02 | 3.38E-02 | tRNA-Leu |
| PA0608 | 3.53E+01 | 3.41E-02 | phosphoglycolate phosphatase |
| PA0265 | 2.96E+01 | 3.54E-02 | glutarate-semialdehyde dehydrogenase DavD |
| PA1798 | 2.99E-02 | 3.56E-02 | two-component sensor ParS |
| PA1383 | 3.09E+01 | 3.59E-02 | hypothetical protein PA1383 |
| PA4679 | 8.63E-03 | 3.60E-02 | hypothetical protein PA4679 |
| PA0620 | 6.41E-02 | 3.68E-02 | bacteriophage protein |
| PA4499 | 1.32E+01 | 3.72E-02 | transcriptional regulator |
| PA4371 | 1.02E+02 | 3.85E-02 | hypothetical protein PA4371 |
| PA2010 | 2.22E+01 | 4.13E-02 | transcriptional regulator |
| PA3298 | 2.42E+01 | 4.25E-02 | hypothetical protein PA3298 |
| PA1278 | 1.15E+01 | 4.37E-02 | adenosylcobinamide kinase/adenosylcobinamide-phosphate guanylyltransferase |
| PA3572 | 3.02E+01 | 4.82E-02 | hypothetical protein PA3572 |
| PA4659 | 2.09E+01 | 4.82E-02 | transcriptional regulator |
| PA1796.4 | 1.21E+01 | 4.82E-02 | tRNA-His |
| PA2699 | 8.39E-02 | 4.82E-02 | hypothetical protein PA2699 |

**Figure 4.**
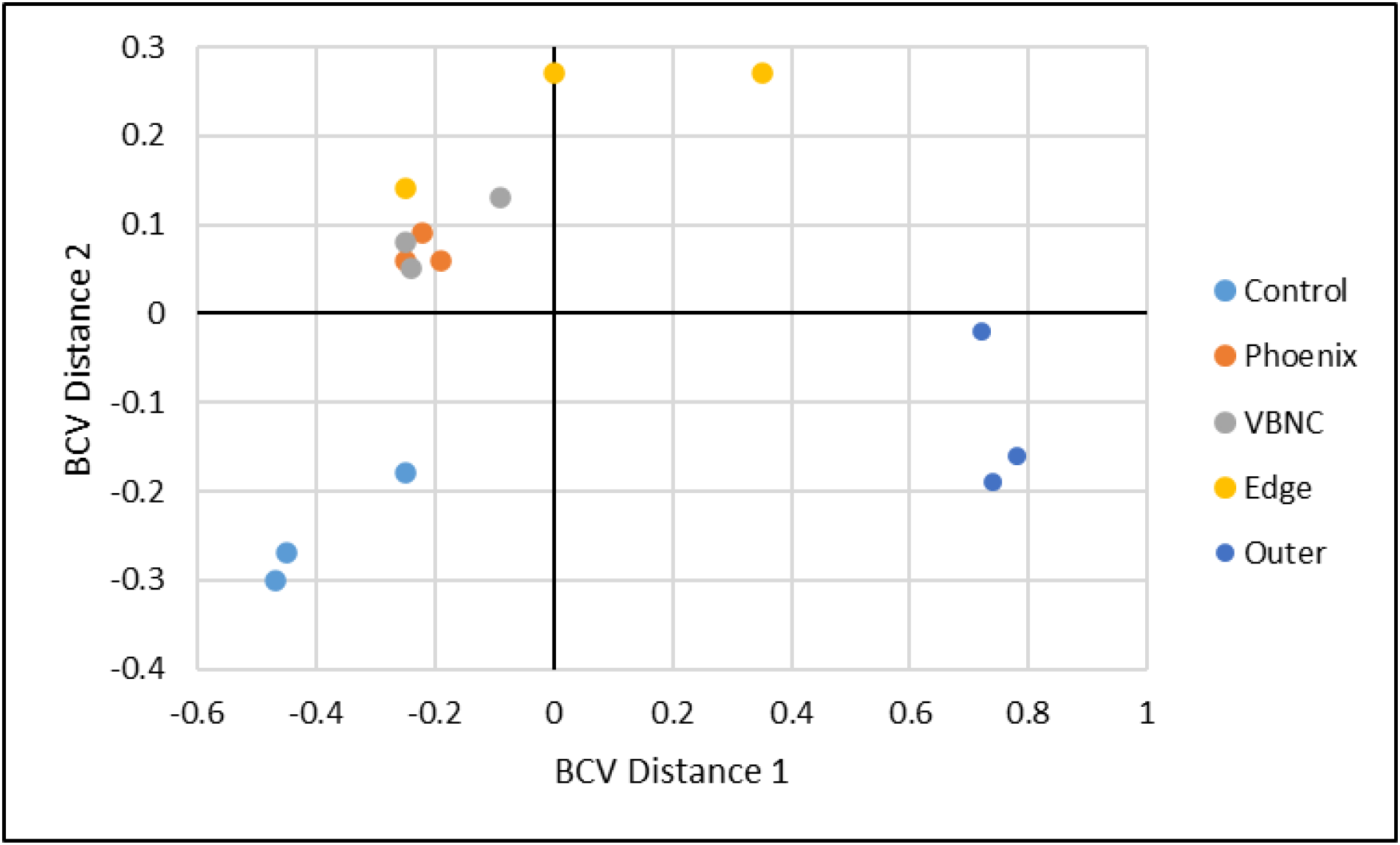
Multi-dimensional scaling (MDS) plot showing clustering of isolates. An MDS plot was generated using biological coefficients of variation (BCV) from transcript counts for each isolate sample. Samples from each group cluster well together aside from the Edge samples (yellow), and the phoenix colonies and VBNC-like colonies overlap considerably. Control lawn, edge, and outer samples cluster on their own and separately from the phoenix and VBNC-like samples.

**Figure 5.**
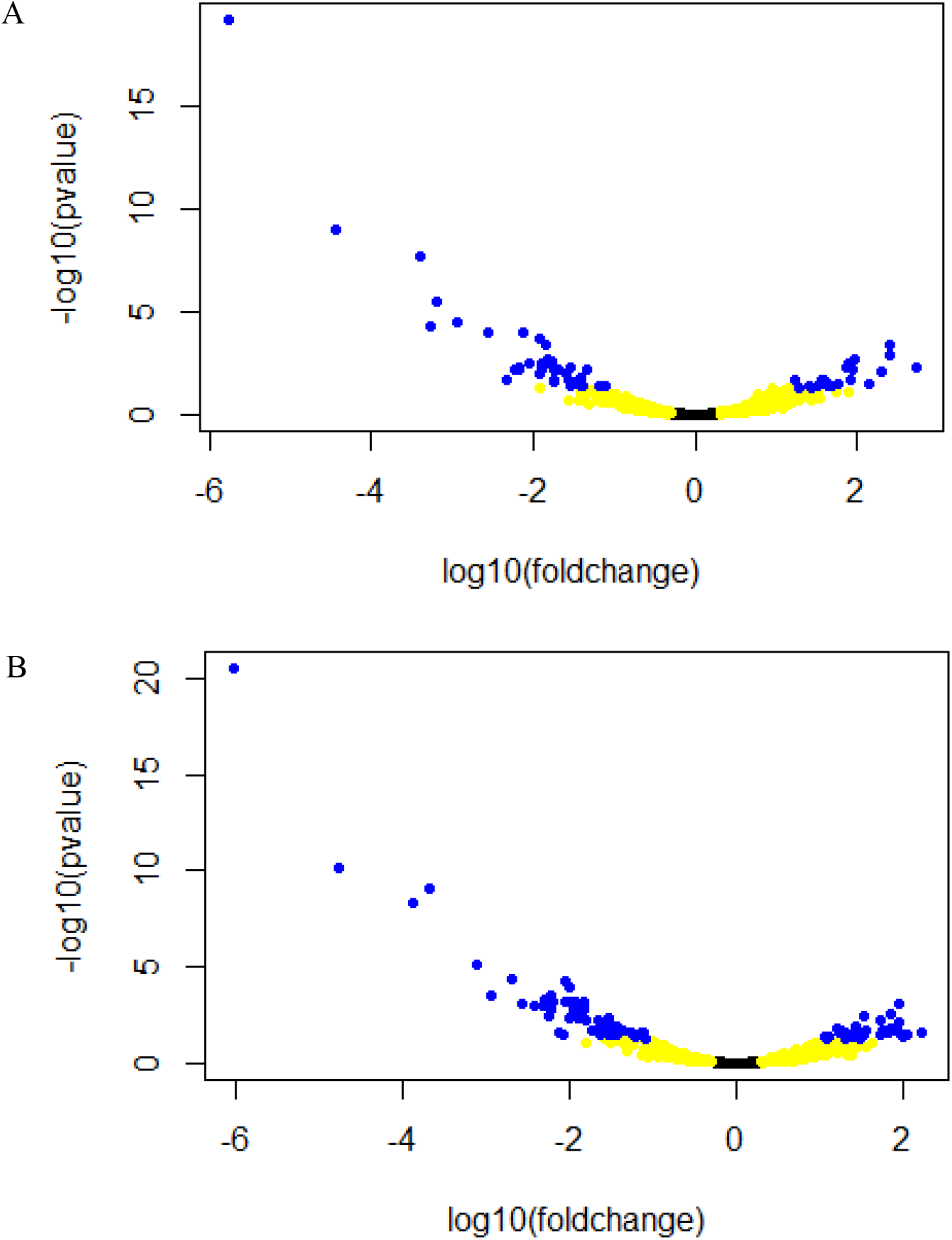
Volcano plots of DEGs. A. Comparison of phoenix colony transcript counts to control lawn transcript counts showed 63 DEGs. B. Comparison of VBNC-like colony transcript counts to control lawn transcript counts showed 90 DEGs. A-B. Yellow dots indicate genes with at least a two-fold change. Red dots indicate a corrected p-value of 0.05 or less. Blue dots indicate both at least a two-fold change and corrected p-value less than 0.05. Black dots indicate genes with no-significant difference in transcription.

### Phoenix Colony Emergence is Eliminated by Disruption of PA3626

Transposon mutants, from the Colin Manoil (15) *P. aeruginosa* transposon mutant library, for each DEG were screened for changes in the frequency of phoenix colony emergence. Nine DEGs did not have a corresponding transposon mutant available in the library for screening (Table 3). Yet, the transposon mutant for PA3626 produced a complete lack of phoenix colony emergence when exposed to tobramycin (n=6). PA3626 encodes for the pseudouridine synthase D (TruD) (16). Complementation of the full PA3626 gene into the PA3626 transposon mutant, using a pUCP18:PA3626 plasmid, restored the frequency of phoenix colony formation to wild-type levels (Figure 6), indicating the importance of TruD in phoenix colony emergence.

**Table 3.** Phoenix colony DEGs remaining after transposon mutant screen. After screening DEG associated transposon mutants, PA3626 was found to have a complete loss of phoenix colony emergence in the presence of tobramycin (n=6). The remaining genes in this table had no available transposon mutant for screening. Red indicates genes which were significantly upregulated and blue indicates genes which were significantly downregulated.

| Gene | Fold Change | q-value | Protein Product Annotation |
| --- | --- | --- | --- |
| PA4746 | 2.63E+01 | 4.79E-02 | hypothetical protein PA4746 |
| PA3626 | 3.78E-02 | 4.23E-02 | tRNA pseudouridine synthase D |
| PA2819.3 | 4.88E+01 | 3.79E-02 | tRNA-Glu |
| PA4529 | 5.78E+01 | 3.63E-02 | dephospho-CoA kinase |
| PA4746.2 | 1.42E+02 | 3.63E-02 | tRNA-Leu |
| PA4746.1 | 8.10E+01 | 2.21E-02 | tRNA-Met |
| PA0103a | 1.66E-02 | 3.31E-03 | hypothetical protein PA0103a |
| PA2139 | 1.67E-02 | 2.64E-03 | pseudogene |
| PA4745 | 2.43E+02 | 1.30E-03 | transcription elongation factor NusA |
| PA0526 | 1.68E-06 | 6.45E-20 | hypothetical protein PA0526 |

**Figure 6.**
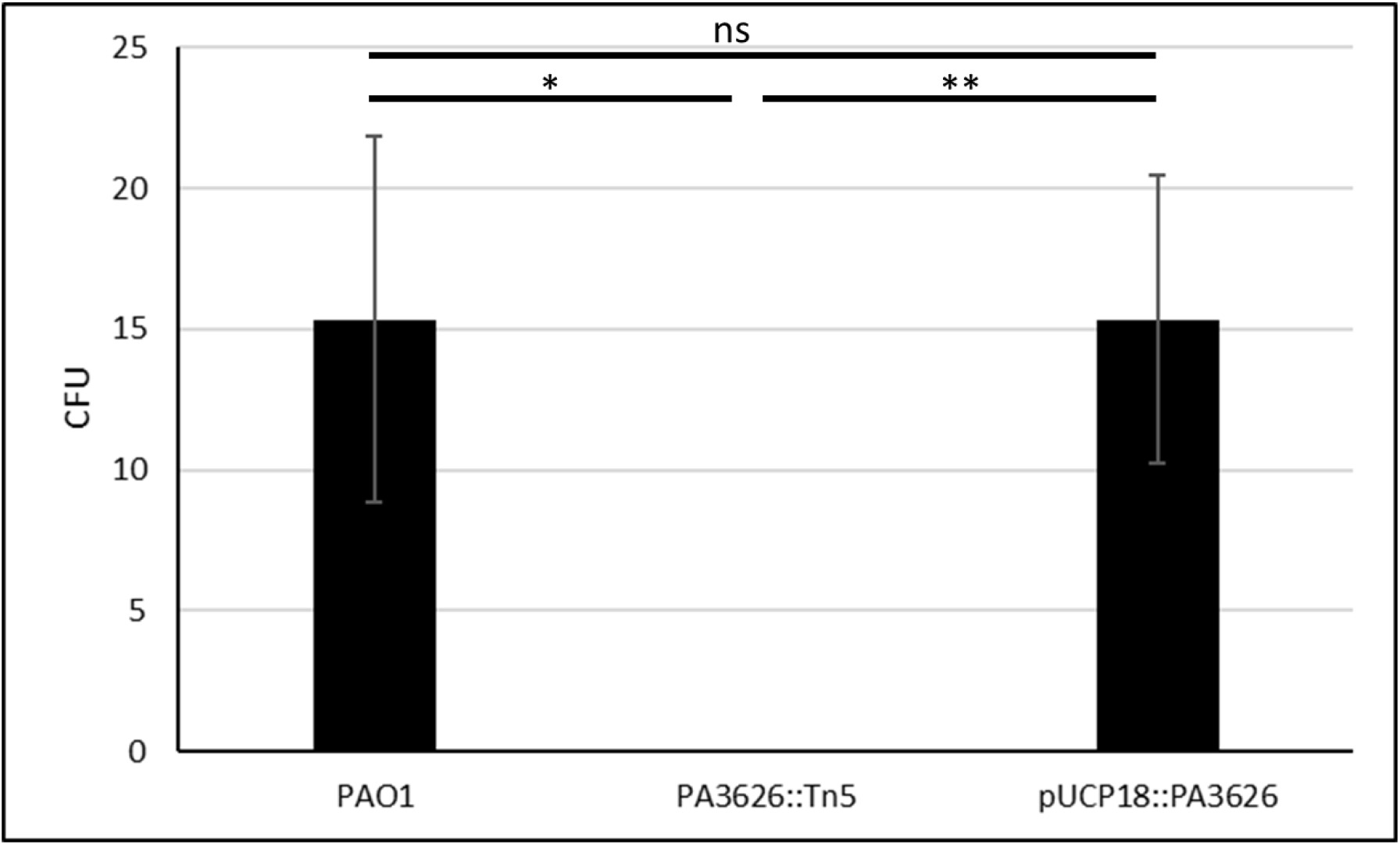
Complementation of PA3626 shows rescue of the phoenix phenotype. Phoenix colony counts were obtained for wild-type *P. aeruginosa* PAO1, the PA3626 transposon mutant, and the PA3626 complementation strain. After complementation of the transposon mutant, phoenix colony counts returned to wild-type levels (p=1), further confirming the importance of the tRNA pseudouridine synthase D in the emergence of phoenix colonies. *p<0.05, **p<0.01, n=3.

## Discussion

*P. aeruginosa* is an important bacterial pathogen implicated in both superficial and life-threatening infections that has readily developed tolerance and resistance to multiple antibacterial drugs. In this study, both the genomes and transcriptomes of both phoenix colonies and VBNC-like colonies were examined. Whole genome sequencing with pooling of the samples was used to identify conserved driver mutations. While VBNC-like isolates display a phenotype different than that of wild-type *P. aeruginosa*, both phenotypes were genotypically identical. However, when compared to wild-type *P. aeruginosa*, phoenix colonies have a SNP in the PA4673 gene which is predicted to encode a GTP-binding protein similar to YchF in *S. pombe* (13, 14). This SNP leads to a modification of the protein structure in an external region of the gene and could possibly lead to a change in a binding site. GTP binding proteins, also known as G proteins, are molecular switches which hydrolyze GTP to GDP and are bound to the membrane to allow for downstream effects to be stimulated by extracellular stimuli (17). A modification in the ability of a G protein to bind to a substrate may have a major impact to downstream effectors within the pathway and may allow phoenix colonies to transiently tolerate the presence of high concentrations of antibiotic.

Transcriptionally, both phoenix colonies and VBNC-like colonies were found to cluster separately from cells in the control lawn biofilm which had not been exposed to antibiotics, the edge of the ZOC, and the outer background lawn controls, while clustering well with each other. It is interesting that both the phoenix colonies and VBNC-like colonies are transcriptomically similar to each other despite appearing as two distinct phenotypes. It is possible that VBNC-like colonies could be a subset of phoenix colonies which lack the ability to be cultured due to a dependence on a metabolite in the initial environment from either the original biofilm lawn or surrounding variant colonies. It is also interesting to note that no significant metabolic pathway changes were found in phoenix colonies or VBNC-like colonies which could account for the antibiotic tolerance exhibited by these phenotypes. This contrasts the typical tolerance phenotypes found in *P. aeruginosa* including persister cells, small colony variants, and adaptive resistance colonies, which enter a state of dormancy, have growth defects, or are otherwise metabolically driven (10, 18–22).

Upon further examination of the DEGs associated with phoenix colonies, one gene was found that when screened showed a lack of phoenix colony emergence: PA3626, which encodes for tRNA pseudouridine synthase D (TruD). Tobramycin, the antibiotic associated with the emergence of phoenix colonies (9), is an aminoglycoside. Aminoglycosides bind irreversibly to the 30S portion of the bacterial ribosome, leading to mismatching binding of tRNA to the mRNA due to a steric hindrance during translation (23). Antibiotic resistance to aminoglycosides can be conferred through modification of the ribosomal target site (24). Additionally, it has been shown that aminoglycosides also interact directly with tRNA and can lead to a loss of interaction between the T- and D- loops as well as unfolding of the D-stem (25, 26). TruD converts uracil-13 in the D-loop to pseudouridine which may influence the way that aminoglycosides are able to interact with the tRNA (27). Knockout of TruD by transposon mutagenesis led to an elimination of phoenix colony emergence which was able to be rescued to wild-type levels by plasmid complementation. This finding is interesting as it leads us to believe that, in addition to the ribosome, tobramycin may also be interacting with tRNA leading to cell death. Phoenix colonies may then be modifying their tRNA with TruD to increase tRNA stability and allow for survival with exposure to tobramycin, although this is something that will need to be followed up on for confirmation.

Further research, including epigenetic exploration, is needed to come to a complete understanding of both the phoenix colonies and VBNC-like colonies. By understanding how these and other bacterial phenotypes are able to survive in the presence of antibiotics, we will be better prepared to control the rise of antibiotic tolerant and resistant infections.

## Materials and Methods

### Bacterial strains and culture conditions

The bioluminescent strain *P. aeruginosa* Xen41 (Xenogen Corp., USA) was used for the imaging portion of this study (Figure 1). Additionally, *P. aeruginosa* PAO1 (15), which is also the parent strain for *P. aeruginosa* Xen41, was used in this study. Stock culture plates were prepared by streaking from glycerol stock cultures which were stored at −80°C onto fresh Luria-Bertani (LB) agar plates. The streaked plates were incubated at 37°C for 24 hours before being examined for individual colony growth and proper morphology. Individual colonies were isolated from stock culture plate and transferred to 20 mL of LB broth. Broth cultures were incubated overnight in a shaking incubator set to 37°C and 200 rpm.

### Preparation of tobramycin loaded calcium sulfate beads

Antibiotic loaded calcium sulfate beads were added to pre-formed biofilm lawns. 240 mg of tobramycin (Sigma-Aldrich) per 20 g of CaSO_4_ hemi-hydrate (Sigma-Aldrich), a ratio commonly used by orthopedic surgeons for treatment of periprosthetic joint infections (PJIs), was used (28). After mixing of the tobramycin and CaSO_4_ powders, sterile water was added and mixed to produce a thick paste. The paste was then spread into silicone molds (Biocomposites Ltd.) to form 4.5 mm diameter, hemispherical beads. The beads were allowed to dry at room temperature overnight before being removed from the mold and stored at 4°C before use.

### Preparation of phoenix and VBNC-like colonies of P. aeruginosa

Phoenix colonies were obtained as per Sindeldecker et al. (9). Briefly, an overnight bacterial culture of *P. aeruginosa* PAO1 was diluted to an OD_600_ of 0.1 using sterile LB broth. One hundred μL of the diluted culture was spread onto LB agar contained in 100 mm diameter, polystyrene plates (Fisher Scientific, USA). The plates were incubated at 37°C with 5% CO_2_ for 24 hours in a humidified incubator (Heracell 150i, Thermo Scientific) to allow a lawn biofilm of *P. aeruginosa* to develop. After the 24 hour lawn biofilm was prepared, a tobramycin-loaded calcium sulfate bead was placed in the center of the plate and pushed into the agar using sterile forceps. The plates were then incubated further at 37°C with 5% CO_2_ for an additional 96 hours. Plates were visually checked daily for the appearance of a zone of clearance (ZOC) as well as colonies emerging within this zone. Additionally, 72 hour lawn biofilms were prepared in the same fashion but without tobramycin beads for use as a control lawn.

### Isolation of Samples to be used for Whole Genome Sequencing

Phoenix colonies, VBNC-like colonies, and wild-type *P. aeruginosa* PAO1 colonies were isolated for whole genome sequencing. In short, phoenix colony plates were generated as above. After the emergence of the variant colonies on three replicate plates, 96 colonies from each plate were isolated into individual aliquots of 100 μL of PBS and stored at −20°C. Before placing the colony isolate samples at −20°C, 5 μL was used to inoculate 200 μL of LB broth containing 5 μg/mL of tobramycin in a well of a 96-well plate (Corning, Sigma-Aldrich). An additional 5 μL was added to a corresponding well of another 96-well plate containing 200 μL of LB broth. The plates were then incubated for 96 hours at 37°C with 5% CO_2_. After incubation, turbidity in the wells of each plate was visually compared. Variants which showed a lack of growth in the LB broth as well as the corresponding well of LB broth containing the tobramycin were defined to be VBNC-like colonies. Phoenix colonies were defined as those showing a lack of growth in the LB broth containing tobramycin while having growth in the LB only wells. An overnight broth culture of *P. aeruginosa* PAO1 was also grown as above to provide a wild-type sample for DNA extraction.

### Determination of Variant Colony Antibiotic Susceptibility

Isolates which grew from the previous section were subjected to a Kirby-Bauer assay to determine whether they were resistant or susceptible to tobramycin. Each isolate was diluted to an OD_600_ of 0.1. 100 μL of each diluted culture was spread onto sterile LB agar plates. A sterile, filter paper disk was placed in the center of each plate and 10 μg of tobramycin was placed on each disk. These plates were incubated at 37°C with 5% CO_2_ for 24 hours. After incubation, the diameter of the zone of inhibition produced was measured and compared to the Clinical and Laboratory Standards Institute (CLSI) guidelines to determine if they were resistant or susceptible (phoenix colonies). Four phoenix colonies and four VBNC-like colonies were randomly selected to be used in RNA extraction.

### Whole Genome Sequencing

DNA extraction was performed for each of the above isolated samples by following manufacturer’s instructions using the GenElute™ Bacterial Genomic DNA kit (Sigma-Aldrich). After gDNA extraction, samples were pooled to obtain samples of 400 ng gDNA to identify conserved driver mutations. Library preparation and barcoding was completed using the Rapid Barcoding Kit (Nanopore). A Nanopore MinION Sequencer was then used to complete bacterial whole genome sequencing using default settings. After sequencing was completed, quality control was performed using the software Epi2ME (Nanopore). Reads were then trimmed using Porechop (29) and aligned to a reference *P. aeruginosa* PAO1 genome (GCF_000006765.1, available for download from the Pseudomonas Genome database, (30)) using graphmap (31). Reads were sorted and indexed using samtools (32). Structural variants were identified using sniffles (33) and SNPs were identified using BCFtools (34).

### Isolation of Samples to be used for RNAseq

Once phoenix colony plates had been prepared, samples were taken for use in RNA sequencing. Seventy-two hours post tobramycin bead placement, triplicate samples were taken from the edge of the zone of clearance (ZOC) where phoenix colonies would be visible on the following day. Ninety-six hours post tobramycin bead placement, triplicate samples from the outer background lawn were taken and variant colonies which had emerged in the ZOC were isolated for phenotype characterization (phoenix colonies, VBNC-like colonies, or resistant mutants). Additionally, triplicate samples were taken from bacterial lawns grown for 72 hours that had not been exposed to antibiotics. All samples and isolates were placed into 100 μL of RNAlater (Ambion) and stored at −20°C to protect the RNA from degradation until needed for RNA extraction. A control study was done in which 40 colonies of wild-type *P. aeruginosa* PAO1 were isolated into LB broth and 40 colonies were isolated into RNAlater before being transferred to LB broth. No significant difference was seen in viability of the colonies exposed to RNAlater (p=1.0). For the variant colony isolates, immediately after being mixed in the RNAlater, a 5 μL sample was taken and used to inoculate 15 mL of sterile LB broth. These broth cultures were incubated at 37°C with 200 rpm shaking overnight. Cultures which did not grow in the overnight broth cultures were deemed to be VBNC-like colonies. The cultures which did grow were then further examined to determine their level of susceptibility to tobramycin.

### RNA Sequencing

RNA extraction was performed for each sample using the Direct-zol RNA Miniprep Kit (Zymo) following manufacturer’s instructions. RNA quantity was measured using Qubit 3.0. In order to remove ribosomal RNA from isolated samples, the RiboZero Bacteria Kit (Illumina) was used as per manufacturer’s instructions. Barcoded RNAseq libraries were then generated using the ScriptSeq™ v2 Kit (Illumina) per the manufacturer’s instructions. Samples were sequenced using paired-end (2 x 150 bp) sequencing on an Illumina HiSeq 4000 device. Sequencing reads were quality controlled using FastQC (35) and mapped to the reference genome for *P. aeruginosa* PAO1 (GCF_000006765.1, available for download from the Pseudomonas Genome database, (30)) using STAR (36). After read mapping, sequences were visualized to ensure adequate genome coverage using IGV (v2.4.13, (37)). Transcript counts were obtained using HTseq (38) and were further analyzed for differential gene expression using the R package edgeR (v3.24.3, (39)). Multi-dimensional scaling plots were also generated using the plotMDS function within edgeR. Volcano plots were generated using the plot function in R. Genes found to be differentially expressed when comparing phoenix colony and VBNC-like colony samples to 72 hour unexposed lawn samples were submitted to DAVID 6.8 (40, 41), KEGG (42), and Gene Ontology (43, 44) for analysis.

### Screening DEG Transposon Mutants

Transposon mutants (obtained from the lab of Dr. Colin Manoil, (15)) corresponding to each of the identified differentially expressed genes were screened in triplicate replicates for their ability to produce phoenix colonies. A biofilm lawn was generated for each mutant, a tobramycin loaded CaSO_4_ bead was placed, and phoenix colonies were generated as described above. Upon phoenix colony emergence, phoenix colonies were distinguished from resistant colonies and VBNC-like colonies using replica plating as per Sindeldecker et al. (9). Briefly, a sterile, cotton velveteen square (150 × 150 mm) was draped over a PVC replica plating block and held in place using an aluminum replica plater ring. The tobramycin loaded CaSO_4_ bead was removed from the plate using a sterile, plastic loop. The plate was marked to indicate a 12 o’clock position and gently placed on the velveteen square. The plate was then gently tapped down before being removed from the replica plater. A fresh, sterile LB agar plate containing 5 μg/mL of tobramycin and marked at the 12 o’clock position was placed on the velveteen square and tapped down in the same fashion before being removed. A fresh, sterile LB agar plate marked at the 12 o’clock position was placed on the velveteen square and tapped down before being removed. Replica plates were incubated for 24 hours at 37°C with 5% CO_2_. After incubation, the colony pattern on the plates was compared and colonies which appeared on the LB agar replica plate but not on the LB agar replica plate containing tobramycin were determined to be phoenix colonies. Transposon mutant phoenix colony counts were obtained and compared to control *P. aeruginosa* PAO1 counts.

### Creation of a PA3626 Complementation Plasmid

*P. aeruginosa* PAO1 gDNA was extracted using the GenElute™ Bacterial Genomic DNA kit. After extraction, PCR was performed to amplify the PA3626 gene using the manufacturer’s protocols for Phusion Taq polymerase (New England Biolabs) and forward (5’-GCGCGGATCCATGAGCGTTCTCGGCGAA) and reverse (5’-GCGCTCTAGATCAGTATGCGCATGGGTT) primers (Integrated DNA Technologies). After amplification, the PA3626 product was run on an agarose gel and extracted using the GenElute™ Gel Extraction kit. A DNA digestion was then completed using the restriction enzymes BamHI and XbaI (New England Biolabs) on both the PA3626 product and pUCP18 plasmid following the New England Biolabs protocol. DNA ligation was then completed using manufacturer’s protocol for the T4 DNA ligase (Thermo Fisher) and a 7:1 ratio of insert to plasmid. After ligation, the plasmid was electroporated into *P. aeruginosa* PAO1 PA3626∷Tn5. Briefly, 1 mL of an overnight culture of PAO1 PA3626∷Tn5 was centrifuged at max speed in a microcentrifuge for 1 min. The pellet was then washed three times in 10% sucrose in sterile water before being resuspended in 100 μL of 10% sucrose. 5 μL of the ligated plasmid was added and the mixture was transferred to a 2 mm electroporation cuvette. The cuvette was then electroporated at 25 μF, 200 Ω, and 2.5 kV. 1 mL of sterile LB was then added to the cuvette and the culture was transferred to a 1.5 mL tube and incubated at 37°C with 200 rpm shaking for 1 hr. After incubation, the culture was spread on LB agar containing 100 μg/mL of carbenicillin and incubated overnight at 37°C with 5% CO_2_. Isolates were then cultured and plasmids were extracted using the Qiagen Spin Miniprep kit. Plasmids were then digested using BamHI and XbaI restriction enzymes, and an agarose gel was run to confirm the appropriately sized bands were present.

### Screening Complementation Strain for Phoenix Colony Emergence

The PA3626 complementation strain was plated and exposed to 1 mg of tobramycin as above in order to allow variant colonies to emerge. After variant colony emergence, the plates were replica plated onto fresh LB agar and LB agar containing 5 μg/mL of tobramycin. Replica plates were incubated for 24 hours at 37°C with 5% CO_2_. After incubation, plates were compared to each other and colonies which grew on the original plate and LB only replica plate but did not grow on the replica plate containing tobramycin were counted as phoenix colonies.

### Protein Modeling

Amino acid sequences for wild-type PA4673 and PA4673 containing the identified SNP were submitted for protein structure and functional modeling to Phyre2 on an internal development server (12) using the intensive modeling mode. Models were visualized using RasMol (45). In addition, amino acid sequences for both wild-type and mutant PA4673 were submitted to the ExPASy server (46) to examine changes in hydrophobicity caused by the predicted structural changes in the mutant protein.

### Statistical analysis

All experiments were performed with a minimum of triplicate technical replicates. ANOVA analysis was completed in edgeR for differentially expressed genes with significance set at a q-value of 0.05. ANOVA analysis was completed by GraphPad Prism (v8.2.1) for all other studies with a p-value of 0.05 being considered significant.

### Bioluminescence Imaging

Bioluminescence imaging was performed using an *in vitro* imaging system (IVIS). Plates were exposed for thirty second exposures to obtain images. A pseudo-color heatmap was then applied to aid in visualization.

## Acknowledgements

This work was supported in part by the Ohio State University College of Medicine and R01 NIH- GM124436 (PS), R01AI148788 (MZA), an NSF CAREER Award 2046863 (MZA), the American Heart Association grant AHA 20PRE35200201 (MJD), NIH GM132254 (JDA), and NIH GM084065-11 (JDA). We would like to thank the Genomics Shared Resource of the Ohio State University Comprehensive Cancer Center for performing the RNA sequencing. We would also like to thank Robert Fillinger for technical assistance with the RNAseq data analysis. Additionally, we would like to thank the lab of Dr. Daniel Wozniak for providing strains from their stocks of Dr. Colin Manoil’s *P. aeruginosa* transposon library.

## References

1. Hoiby N, Ciofu O, Bjarnsholt T. 2010. Pseudomonas aeruginosa biofilms in cystic fibrosis. Future Microbiol 5:1663–74.

2. Serra R, Grande R, Butrico L, Rossi A, Settimio UF, Caroleo B, Amato B, Gallelli L, de Franciscis S. 2015. Chronic wound infections: the role of Pseudomonas aeruginosa and Staphylococcus aureus. Expert Rev Anti Infect Ther 13:605–13.

3. Shah NB, Osmon DR, Steckelberg JM, Sierra RJ, Walker RC, Tande AJ, Berbari EF. 2016. Pseudomonas Prosthetic Joint Infections: A Review of 102 Episodes. J Bone Jt Infect 1:25–30.

4. Lewis K. 2008. Multidrug tolerance of biofilms and persister cells. Curr Top Microbiol Immunol 322:107–31.

5. Zavascki AP, Carvalhaes CG, Picao RC, Gales AC. 2010. Multidrug-resistant Pseudomonas aeruginosa and Acinetobacter baumannii: resistance mechanisms and implications for therapy. Expert Rev Anti Infect Ther 8:71–93.

6. Hall-Stoodley L, Stoodley P. 2005. Biofilm formation and dispersal and the transmission of human pathogens. Trends Microbiol 13:7–10.

7. Lewis K. 2010. Persister cells. Annu Rev Microbiol 64:357–72.

8. Oliver A, Canton R, Campo P, Baquero F, Blazquez J. 2000. High frequency of hypermutable Pseudomonas aeruginosa in cystic fibrosis lung infection. Science 288:1251–4.

9. Sindeldecker D, Moore K, Li A, Wozniak DJ, Anderson M, Dusane DH, Stoodley P. 2020. Novel Aminoglycoside-Tolerant Phoenix Colony Variants of Pseudomonas aeruginosa. Antimicrob Agents Chemother 64.

10. Brauner A, Fridman O, Gefen O, Balaban NQ. 2016. Distinguishing between resistance, tolerance and persistence to antibiotic treatment. Nat Rev Microbiol 14:320–30.

11. Ramamurthy T, Ghosh A, Pazhani GP, Shinoda S. 2014. Current Perspectives on Viable but Non-Culturable (VBNC) Pathogenic Bacteria. Front Public Health 2:103.

12. Kelley LA, Mezulis S, Yates CM, Wass MN, Sternberg MJ. 2015. The Phyre2 web portal for protein modeling, prediction and analysis. Nat Protoc 10:845–58.

13. R.K. Kniewel JAB, C.D. Lima. Structure of the S. pombe YchF GTP-binding protein.

14. Teplyakov A, Obmolova G, Chu SY, Toedt J, Eisenstein E, Howard AJ, Gilliland GL. 2003. Crystal structure of the YchF protein reveals binding sites for GTP and nucleic acid. J Bacteriol 185:4031–7.

15. Jacobs MA, Alwood A, Thaipisuttikul I, Spencer D, Haugen E, Ernst S, Will O, Kaul R, Raymond C, Levy R, Chun-Rong L, Guenthner D, Bovee D, Olson MV, Manoil C. 2003. Comprehensive transposon mutant library of Pseudomonas aeruginosa. Proc Natl Acad Sci U S A 100:14339–44.

16. Kaya Y, Ofengand J. 2003. A novel unanticipated type of pseudouridine synthase with homologs in bacteria, archaea, and eukarya. RNA 9:711–21.

17. Rosenbaum DM, Rasmussen SG, Kobilka BK. 2009. The structure and function of G-protein-coupled receptors. Nature 459:356–63.

18. Lewis K. 2007. Persister cells, dormancy and infectious disease. Nat Rev Microbiol 5:48–56.

19. Renbarger TL, Baker JM, Sattley WM. 2017. Slow and steady wins the race: an examination of bacterial persistence. AIMS Microbiol 3:171–185.

20. Haussler S, Tummler B, Weissbrodt H, Rohde M, Steinmetz I. 1999. Small-colony variants of Pseudomonas aeruginosa in cystic fibrosis. Clin Infect Dis 29:621–5.

21. Schiessl KT, Hu F, Jo J, Nazia SZ, Wang B, Price-Whelan A, Min W, Dietrich LEP. 2019. Phenazine production promotes antibiotic tolerance and metabolic heterogeneity in Pseudomonas aeruginosa biofilms. Nat Commun 10:762.

22. McCurtain JL, Gilbertsen AJ, Evert C, Williams BJ, Hunter RC. 2019. Agmatine accumulation by Pseudomonas aeruginosa clinical isolates confers antibiotic tolerance and dampens host inflammation. J Med Microbiol 68:446–455.

23. Jana S, Deb JK. 2006. Molecular understanding of aminoglycoside action and resistance. Appl Microbiol Biotechnol 70:140–50.

24. Bryan LE, Haraphongse R, Van den Elzen HM. 1976. Gentamicin resistance in clinical-isolates of Pseudomonas aeruginosa associated with diminished gentamicin accumulation and no detectable enzymatic modification. J Antibiot (Tokyo) 29:743–53.

25. Walter F, Putz J, Giege R, Westhof E. 2002. Binding of tobramycin leads to conformational changes in yeast tRNA(Asp) and inhibition of aminoacylation. EMBO J 21:760–8.

26. Wang B, Wilkinson KA, Weeks KM. 2008. Complex ligand-induced conformational changes in tRNA(Asp) revealed by single-nucleotide resolution SHAPE chemistry. Biochemistry 47:3454–61.

27. Kaya Y, Del Campo M, Ofengand J, Malhotra A. 2004. Crystal structure of TruD, a novel pseudouridine synthase with a new protein fold. J Biol Chem 279:18107–10.

28. Howlin RP, Brayford MJ, Webb JS, Cooper JJ, Aiken SS, Stoodley P. 2015. Antibiotic-loaded synthetic calcium sulfate beads for prevention of bacterial colonization and biofilm formation in periprosthetic infections. Antimicrob Agents Chemother 59:111–20.

29. Wick R. 2018. Porechop. https://github.com/rrwick/Porechop. Accessed

30. Winsor GL, Griffiths EJ, Lo R, Dhillon BK, Shay JA, Brinkman FS. 2016. Enhanced annotations and features for comparing thousands of Pseudomonas genomes in the Pseudomonas genome database. Nucleic Acids Res 44:D646–53.

31. Sovic I, Sikic M, Wilm A, Fenlon SN, Chen S, Nagarajan N. 2016. Fast and sensitive mapping of nanopore sequencing reads with GraphMap. Nat Commun 7:11307.

32. Li H, Handsaker B, Wysoker A, Fennell T, Ruan J, Homer N, Marth G, Abecasis G, Durbin R, Genome Project Data Processing S. 2009. The Sequence Alignment/Map format and SAMtools. Bioinformatics 25:2078–9.

33. Sedlazeck FJ, Rescheneder P, Smolka M, Fang H, Nattestad M, von Haeseler A, Schatz MC. 2018. Accurate detection of complex structural variations using single-molecule sequencing. Nat Methods 15:461–468.

34. Li H. 2011. A statistical framework for SNP calling, mutation discovery, association mapping and population genetical parameter estimation from sequencing data. Bioinformatics 27:2987–93.

35. Andrews S. FastQC. https://www.bioinformatics.babraham.ac.uk/projects/fastqc/. Accessed

36. Dobin A, Davis CA, Schlesinger F, Drenkow J, Zaleski C, Jha S, Batut P, Chaisson M, Gingeras TR. 2013. STAR: ultrafast universal RNA-seq aligner. Bioinformatics 29:15–21.

37. Robinson JT, Thorvaldsdottir H, Winckler W, Guttman M, Lander ES, Getz G, Mesirov JP. 2011. Integrative genomics viewer. Nat Biotechnol 29:24–6.

38. Anders S, Pyl PT, Huber W. 2015. HTSeq--a Python framework to work with high-throughput sequencing data. Bioinformatics 31:166–9.

39. Robinson MD, McCarthy DJ, Smyth GK. 2010. edgeR: a Bioconductor package for differential expression analysis of digital gene expression data. Bioinformatics 26:139–40.

40. Huang da W, Sherman BT, Lempicki RA. 2009. Systematic and integrative analysis of large gene lists using DAVID bioinformatics resources. Nat Protoc 4:44–57.

41. Huang da W, Sherman BT, Lempicki RA. 2009. Bioinformatics enrichment tools: paths toward the comprehensive functional analysis of large gene lists. Nucleic Acids Res 37:1–13.

42. Kanehisa M, Sato Y. 2020. KEGG Mapper for inferring cellular functions from protein sequences. Protein Sci 29:28–35.

43. Ashburner M, Ball CA, Blake JA, Botstein D, Butler H, Cherry JM, Davis AP, Dolinski K, Dwight SS, Eppig JT, Harris MA, Hill DP, Issel-Tarver L, Kasarskis A, Lewis S, Matese JC, Richardson JE, Ringwald M, Rubin GM, Sherlock G. 2000. Gene ontology: tool for the unification of biology. The Gene Ontology Consortium. Nat Genet 25:25–9.

44. Gene Ontology C. 2021. The Gene Ontology resource: enriching a GOld mine. Nucleic Acids Res 49:D325–D334.

45. Sayle RA, Milner-White EJ. 1995. RASMOL: biomolecular graphics for all. Trends Biochem Sci 20:374.

46. Gasteiger EH, C.; Gattiker, A.; Duvaud, S.; Wilkins, M.R.; Appel, R.D.; Bairoch, A. 2005. Protein Identification and Analysis Tools on the ExPASy Server. Humana Press.

